# Automated detection of blink reflexes evoked by optogenetic stimulation of TRPV1-expressing corneal nociceptors in transgenic mice

**DOI:** 10.64898/2026.06.18.733051

**Authors:** Ki-Soo Jeong, Matthew T. McPheeters, Anagha Chandrasekharan, Ian Beeck, Ananya Veerubhotla, Aditya Roy, Eric Y. Lu, Samer Ghosn, Michael W. Jenkins, Carl Y. Saab

## Abstract

**Background:** Conventional rodent models for the study of corneal pain commonly evoke eye blink reflex using methods that indiscriminately activate polymodal nociceptors, mechanoreceptors, and thermoreceptors at temporal resolutions that don’t closely match the sub-second timescale of underlying neural dynamics.

**New method:** We introduce a novel automated behavioral paradigm for detecting blink reflexes in transgenic TRPV1-ChR2-EYFP mice, enabled by cell-type-specific, millisecond-precision optogenetic stimulation of corneal nociceptors (490 nm light). Using multi-feature quantification, we achieve robust automated detection using univariate and multivariate classifiers.

**Results:** TRPV1-ChR2-EYFP mice exhibited blink reflexes to high-intensity blue light (490 nm, 10 ms pulses) in a threshold-dependent manner (N=3). Blink probability was 77.1 ± 17.1% at high intensity (2.77 mW/mm^2^) versus 4.2 ± 4.2% at low intensity (0.46 mW/mm^2^). Red light (638 nm) produced no intensity-dependent change. Noxious air puff evoked blinks in >95% of trials under all conditions. DeepLabCut-based pose estimation extracted six features quantifying the blink reflex, enabling automated detection with ≥98% accuracy using univariate and multivariate classifiers.

**Comparison with existing methods:** Unlike conventional air puff paradigms, this optogenetic approach enables precise, cell-type-specific stimulation of corneal nociceptors, supporting automated analysis of blink responses at sub-second resolution.

**Conclusions:** This video tracking behavioral method using machine learning algorithms that accurately classify blink versus no-blink enables high-throughput and observer-independent empirical assessment of blink reflex, suggestive of corneal pain. Moreover, inducing blink reflex in TRPV1-ChR2 mice using high-intensity blue light also demonstrates nociceptive-specific behavioral responses analogous to somatosensory optogenetically-evoked hindpaw pain in the same animal genotype.

## Introduction

Understanding how corneal nociceptors encode pain is essential for determining how neural circuit dysfunction contributes to ocular surface diseases such as dry-eye disease and for developing targeted therapeutics. An ideal preclinical model of corneal pain would selectively stimulate corneal nociceptors and produce behavioral responses that are robust, rapid, reproducible and objectively scored. Both clinical and preclinical studies frequently use pressurized air puff to evoke blink reflex as a measure of corneal pain, which fulfills these criteria (Krzyzanowska et al., 2011; Spierer et al., 2016; Varolgünes et al., 1999). However, air puff stimulation lacks anatomical specificity: In addition to activating neurons innervating the ocular surface, it may also recruit periocular afferents adjacent to the eye. In addition, blink reflexes are scored ‘manually’ by visual inspection of real-time behavior of recorded videos.

Even within the cornea, mechanical air puff stimulation simultaneously engages multiple nociceptor types. Mechanical stimuli activate both mechano-nociceptors (mediated via Aδ-fibers) and polymodal nociceptors (mediated via Aδ- and C-fibers), obscuring their individual contributions to nociceptive signaling (Fernández-Trillo et al., 2020). Air puff also produces evaporate cooling at the ocular surface; this cooling effect may activate cold-sensitive thermoreceptors and modulate heat-sensitive polymodal nociceptors (Acosta et al., 2001; Mungalsingh et al., 2025). As a result, blink responses evoked by air puff likely reflect the combined activation of mechanical nociceptors, polymodal nociceptors, and thermoreceptors. Because both the mechanical force and thermal change depend on the air pressure and temperature delivered, simply adjusting stimulus parameters alone does not permit the selective engagement of specific subtypes of corneal nociceptors. This concurrent recruitment limits our ability to isolate the individual contributions of each subtype to corneal pain encoding.

Optogenetic stimulation enables selective activation of corneal nociceptors with millisecond temporal resolution, overcoming the limitations of conventional mechanical stimuli. These light-sensitive opsins have been used extensively to modulate neural circuits and behavior in preclinical models of somatic pain, including mice (Adamantidis et al., 2014; Boyden, 2011; Deisseroth, 2015; Gong et al., 2020; Mattis et al., 2012). By focusing specific wavelengths of light on the cornea without illuminating surrounding tissues, we could in theory selectively activate corneal nociceptors in a controlled and reproducible manner. A behavioral assay for self-report of acute nociception in the transgenic TRPV1-ChR2-EYFP mouse model has been characterized for studying somatic hindpaw pain (Black et al., 2023, 2020). Therefore, we predicted that optogenetic stimulation of corneal nociceptors will elicit reliable blink reflex in this same transgenic model.

Behavioral validation establishes that sensory stimuli reliably evoke a measurable response, providing essential functional context for interpreting concurrently acquired neural data. An ideal evoked behavior should be robust (distinct from other behaviors), rapid (low latency following stimulation), and reproducible across repeated trials. The corneal blink reflex fulfills these criteria, as it is unambiguous, exhibits short latency, and demonstrates high trial-to-trial reproducibility (Varolgünes et al., 1999). We hypothesized that optogenetic activation of corneal afferents in TRPV1-ChR2-EYFP mice would evoke a blink reflex in a manner dependent on light intensity and wavelength. Specifically, we predicted that blink responses would only occur when optogenetic stimuli exceeded a corneal activation threshold. As ChR2/H134R (the variant of channelrhodopsin expressed in this mouse line) exhibits peak excitation near 450 nm, mice were stimulated with 490 nm wavelength blue light at the most oblique angle allowed by the stimulation apparatus (Lin, 2011). To assess wavelength-specificity, 638 nm red light (well outside the peak excitation range of channelrhodopsin) was used as an innocuous control. We therefore expected blink reflexes to occur selectively in response to optogenetic stimulation and not to light outside the excitation spectra.

Behavioral validation requires high-fidelity detection and objective scoring of target behavioral responses. Recent advancements in machine learning address these challenges by providing automated, high-throughput behavioral assessments. While ‘manual’ scoring is a traditional approach, it is inherently limited by subjective observer bias and inter-rater variability, which can introduce inconsistencies. Furthermore, manual assessment is a significant bottleneck, increasing the likelihood of human error and substantially extending processing times. Automation can address these issues of consistency across sessions and extended processing time associated with manual assessment. Machine learning has been applied in various domains: in assessing gait and locomotion, tracking social behavior, and in operantly-conditioned behavioral responses (Gharagozloo et al., 2021; Li et al., 2023; Marks et al., 2020; Sheppard et al., 2022). Here, we applied these techniques in two separate steps: pose estimation of the eye during the blink reflex and multivariate models of binary classification of the evoked blink reflex. This automated framework ensured a standardized, reproducible metric that is scalable across large experimental cohorts.

## 1. Materials and methods

### 1.1. Custom-built system for delivering and observing corneal stimulation

The setup to run an automated behavioral assay of evoked blink response consisted of four main components: a computer equipped with a digital controller (National Instruments, PCIe-8381); a chassis (National Instruments, PXIe-1071) and controller system (National Instruments, PXIe-8381) to sync input and output signals; an acquisition board (Open Ephys); and the custom behavioral platform. This latter behavioral platform held the mouse during testing sessions and was built upon an aluminum breadboard (Thorlabs, MB4560/M). Mice were restrained via headpost on an elevated custom 3D-printed platform during the session.

Corneal stimulation was delivered either via an air puff or a 10 ms pulse of light. Mechanical noxious stimulation was delivered via pressure injection (ASI Imaging, MPPI-3). Optical stimulation was delivered via light from either a 490 nm (Sharp Microelectronics, GH04850B2G) or 638 nm (Thorlabs L638P200) laser diode (blue or red light, respectively). Light from either laser was collimated using a collimator (Thorlabs CFC8-A) to ensure a more consistent beam diameter and enable an unobstructed view of the eye during the session.

### 1.2. Transgenic mice

All methods were performed in accordance with the relevant guidelines and regulations, and animal experiments were approved by the Institutional Animal Care and Use Committee at Cleveland Clinic (CCF), Cleveland, Ohio under protocol #3072. Animals were three male TRPV1-ChR2-EYFP mice, aged 3-6 months, bred at the Cleveland Clinic Behavioral Resource Unit (BRU) from commercially available lines: TRPV1-Cre (#17769) and Ai32 (#24109). The progeny (TRPV1-ChR2-EYFP) were housed in groups of two or more until they underwent headpost implantation, at which point they were housed in single cages to prevent detachment.

### 1.3. Headpost implantation

Mice were anesthetized using isoflurane (3-5%) and maintained throughout the surgical operation. Subcutaneous injections of buprenorphine (Ethiqa XR) and lidocaine were administered to induce analgesia during surgery and local anesthesia, respectively. Once fixed into a stereotactic frame (Kopf Instruments), their heads and adjacent surgical areas were shaved and sterilized using three alternating washes of povidone iodine and 70% ethanol. Ophthalmic ointment was applied to the eyes to prevent damage due to the absent blink response. Skin was excised in a circular window to provide access to the skull, which was then cleaned and scored. A 0.9 mm drill-bit was used to drill holes into the skull using the stereotaxic coordinates for the hindlimb representation of the primary somatosensory cortex (S1HL): -0.5 mm caudal, 1.5 lateral relative to bregma, scaled individually to the distance between bregma and lambda. Stainless steel (Fisher Scientific) screws were set into the skull just above the dura. After resection of the anterior muscles, a custom headpost made from stainless steel or titanium and was positioned above the skull. The headpost and screws were adhered to the skull with dental cement (Metabond). Mice were recovered on oxygen and heated until they were bright, alert, and responsive. Behavioral testing began no less than one week following implantation.

### 1.4. Behavioral assay of evoked blink response to light and air puff

Mice were tested for blink reflex while head-restrained with two different noxious stimuli: mechanical air puff delivered via pressure injector (ASI Imaging, MPPI-3); and noxious blue light of 470 nm or 490 nm and control yellow light of 595 nm wavelength or red light of 638 nm wavelength (see Custom-built system for delivering and observing corneal stimulation). Mice underwent up to 128 trials in each session, not exceeding thirty minutes per session. Stimuli were delivered in a pseudo-random order with 3-6 second ISI. A session ended when the mouse completed all the trials. Video recordings of each trial were scored for blink reflex in a binary system into blink or no-blink trials based on whether the upper and lower eyelids made complete contact immediately following stimulation.

Prior to the blink reflex assessment, trained mice were stimulated at various intensities of light to identify the minimum intensity of light required to elicit a consistent (>90%) blink reflex. This was used to determine the range of stimuli applied during sessions. Mice were also habituated to the testing environment days ahead of testing to acclimate them to the testing setup and minimize stress associated with unfamiliar stimuli.

In separate sessions from evoked blink reflex testing, capsaicin or vehicle was applied prior to recording to induce sensitization-associated squinting, increasing variability in the eye-eyelid configuration during behavioral acquisition for robust pose estimation.

### 1.5. Computational Environment

All analyses were performed using DeepLabCut (v3.0.0rc8) within an Anaconda environment (conda v24.9.2). Deep learning computations were implemented with PyTorch v2.5.1+cu121, using CUDA Toolkit v12.1 for GPU acceleration. Code was executed in Python v3.11.13.

### 1.6. Pose estimation of eyelids via DeepLabCut

For each trial, video recordings were cropped for each trial from 500 milliseconds before to 1000 milliseconds after stimulus delivery (optogenetic or mechanical). A small training set of clips across light conditions (490 nm and 630 nm) was identified across each stimulus intensity tested, as well as with and without the application of capsaicin and vehicle solutions to the eye. These clips were selected to include the widest possible array of responses to various experimental and environmental conditions with the minimum data required. From a video training set of twenty-four trials, up to fourteen frames were extracted and manually labeled for the eight key points we identified: one at the outer and inner corners of the eye (two points); one at the peaks of the upper and lower eyelids (two points); and one halfway between the peak and the corners of each eyelid (four points). Custom models were trained across multiple datasets containing varying numbers of labeled frames (from 100 to 300 total frames), compiled from various corneal stimulation methods, including optogenetic light, control light, and mechanical air stimulation; the custom 250-frame model was selected for the lowest root mean squared error (RMSE) in identifying keypoints while demonstrating high precision and recall. Once the model was successfully trained, the remaining clips were run through the model to identify the eight key points across all frames (**Fig. 3A**). The planar coordinates of these key points across all frames for each trial were then consolidated into a single dataset for all trials.

### 1.7. ‘Manual’ scoring of blink reflexes via custom MATLAB script

For each trial, video recordings were cropped from 500 milliseconds before to 1000 milliseconds after stimulus delivery (optogenetic or mechanical). A custom MATLAB script presented each recording to a blinded observer, who manually scored each blink reflex as no response, partial blink (eyes did not fully close), and complete blink (eyes fully closed). Observer-assigned scores were used as ground-truth labels for evaluating blink detection accuracy. Each trial was independently scored by two observers; discrepant scores were reviewed and resolved by reassessment.

### 1.8. Feature generation and binary thresholds via custom Python script

The planar coordinates for all eight key points for each frame in each clip were aggregated for all trials. These coordinates were then used to calculate six total features for each frame: the Euclidean Distance (the direct distance between the peaks of the upper and lower eyelids) (**Fig. 3B**); the Eye Aspect Ratio or “EAR” (the ratio between the average vertical distance between the upper and lower eyelids to the length of the eyelid) (**Fig. 3C**); the Outer and Inner Eye Angles (angle between the outer and inner corners of the eye to the peaks of the upper and lower eyelids, respectively) (**Fig. 3D**); the Eye Area (the total area of the octagon defined by the eight key points) (**Fig. 3E**); and the Eye Velocity (the change in Euclidean Distance from the previous frame) (**Fig. 3F**). To test whether any given feature could accurately reflect blink reflex, a series of binary thresholds were applied from the minimum to the complete range of possible values for each feature. Acceptable threshold values met the following three criteria: ≥95% accuracy for both experimental (noxious 490 nm light) and control (innocuous 630 nm light) at said given threshold; no more than 5% standard error at said given threshold for both conditions; and no more than 5% difference in accuracy between experimental and control conditions. An ideal threshold was identified from these possible values using the highest average accuracy across both conditions. The minimum and maximum values for each feature across all frames for a given trial were compiled into a separate dataset for multivariate analysis and classification.

### 1.9. Multivariate classification using machine learning via AutoML

Minimum and maximum values across all six features (total twelve values) were identified across all trials along with manually identified blink scores (1 = true; 0 = false) for each trial and aggregated into a single dataset (total 767 trials). These features were then tested for multicollinearity, as the importance of any single predictor becomes unstable, making it difficult to isolate a given feature’s independent impact on predictive accuracy. “Outer angle minimum” was selected for having the highest correlation to the true presence of blink reflex. Other features co-linear with outer angle minimum (absolute correlation ≥ 0.4) were removed from the multivariate dataset; only the features (outer angle min; eye aspect ratio max; velocity max; eye area max), primarily maximum values, were included in the final multivariate analysis. [Variance inflation factor (VIF), which quantifies how much the predictive value of a variable is inflated due to collinearity with other variables, was also factored in: lower VIF scores (VIF ≤ 10) are commonly accepted.] This dataset, along with the manually identified blink scores were fed into seven algorithm families: support vector classifiers (SVC); K-Neighbors Classifiers; Gaussian Naïve Bayes classifiers; Extreme Gradient Boosting classifiers; Categorical Boosting classifiers; Logistic Regression classifiers; and Light Gradient Boosting Machine classifiers. The performance of these classifiers was assessed for balanced accuracy, precision, recall, and receiver operating characteristic (ROC). Confusion matrices and ROC curves were acquired for the highest performing model.

### 1.10. Formulation of a capsaicin compound and vehicle control solutions

A 1 mM stock solution of capsaicin (Sigma-Aldrich, PHR1450-1G) was prepared in 200-proof ethanol. To prepare the ophthalmic formulation, 100 μL of stock solution was diluted with 900 μL of Refresh Optive MEGA-3 lubricant (Allergan) to create a 1:10 dilution. Another 10 μL of this intermediary dilution was added to 990 μL of Refresh Optive MEGA-3 lubricant to achieve a final concentration of 1 μM. The resulting solutions were examined for potential precipitation. The capsaicin-containing eye drops were then transferred to a low-adhesion Eppendorf tube, protected from light with aluminum foil, and stored on ice until use. Vehicle eye-drops were prepared with the same steps with 200-proof ethanol as the stock solution.

## 2. Results

### 2.1. Blue light corneal stimulation of TRPV1-ChR2 elicits a blink reflex

We video-recorded TRPV1-ChR2-EYFP mice during corneal stimulation via air puff or light to identify blink responses. TRPV1-ChR2-EYFP Mice (N=3) were head-restrained and exposed to air puff stimulation sufficient to elicit blinking in >95% of trials, along with optogenetic experimental blue light (490 nm; within the peak excitation range of channelrhodopsin) or innocuous control red light (638 nm; outside the peak excitation range for channelrhodopsin) across multiple light or air intensities. Only one wavelength of light was tested at a time for a given experimental session. Within each session, head-restrained mice received up to 128 randomized trials consisting of either a single-intensity air puff (≤350 ms) or one of seven optogenetic light intensities ranging from near-zero to a high intensity sufficient to elicit near-maximal blink responses (>95%) (**Figure 1**). We ensured that the laser-driven light targeted cornea by collimating the light (beam diameter ∼1.4 mm; by comparison, the average mouse cornea is 2.2 mm in diameter). For purposes of our study, only the left eye was stimulated.

**Figure 1.**
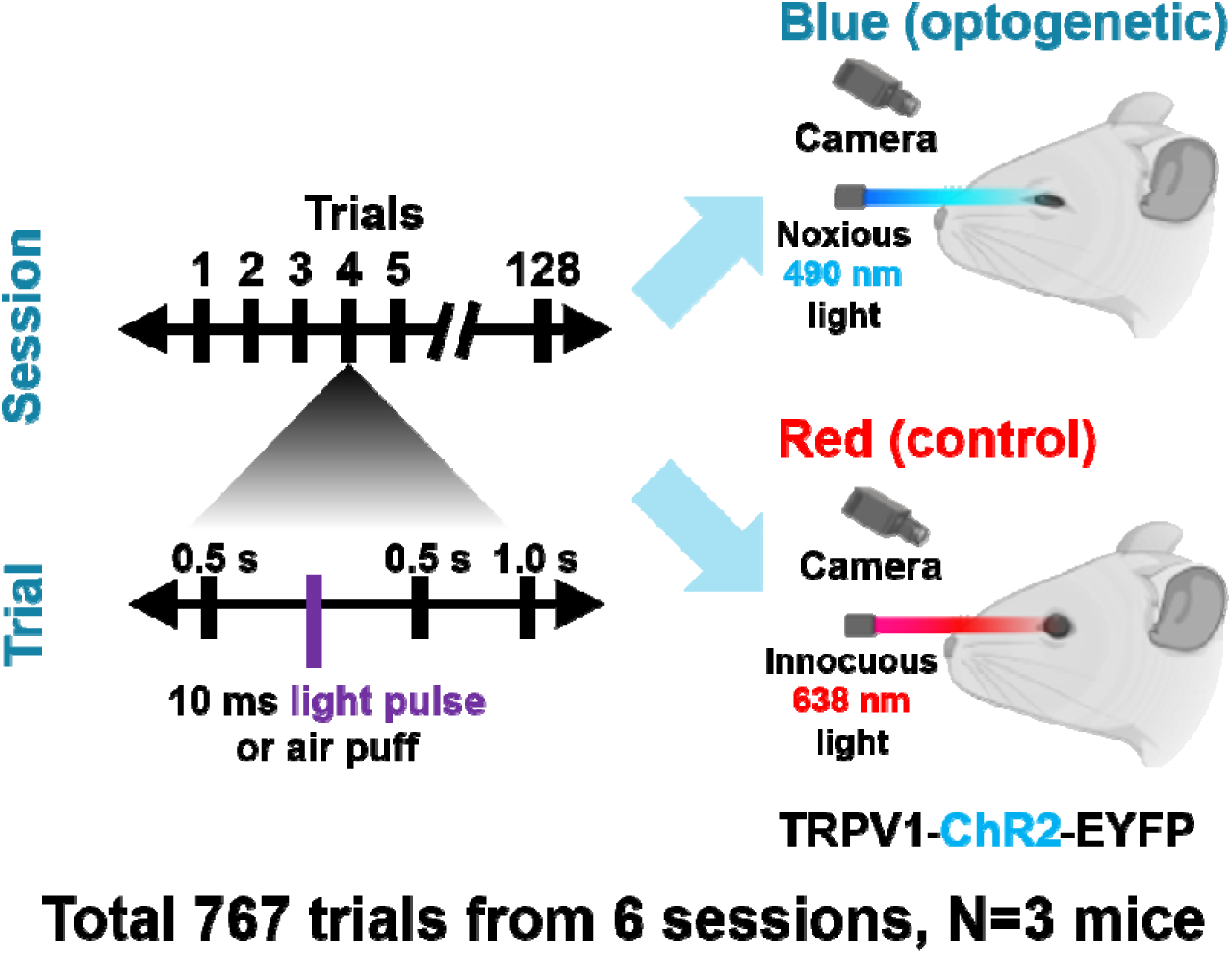
Session-based comparison of blink responses to blue versus red light corneal stimulation in TRPV1-ChR2-EYFP transgenic mice. Each mouse received up to 128 separate, randomized stimuli of light or pressurized air to the cornea and were observed for any subsequent blinking behavior. Mice received either optogenetic blue light (which activates corneal nociceptors) or control red light within a given session; in all sessions, mice also received pressurized air. Trials were defined as the 500 ms before to 1000 ms after stimulus application to the eye.

TRPV1-ChR2-EYFP mice exhibited stimulus-evoked blink reflex in response to optogenetic stimulation with blue light at sufficiently high intensities. At high intensity (2.77 mW/mm^2^), transgenic mice blinked on 77.1 ± 17.1% of trials, significantly greater than the 4.2 ± 4.2% of trials observed with low light intensity (0.46 mW/mm^2^) (**Fig. 2**). By contrast, red light stimulation (638 nm) did not produce a significant change in probability between low (0.26 mW/mm^2^) and high (2.88 mW/mm^2^) intensities, with responses remaining at baseline spontaneous levels (the high intensity reflects the maximum output of the red light system). Noxious air puff stimulation reliably evoked blink responses (>95% across all mice), confirming blink capability at random time intervals. In addition, blink probability scaled with light intensity, consistent with a potential dose-dependent relationship (***Supplementary Figure 2***).

**Figure 2.**
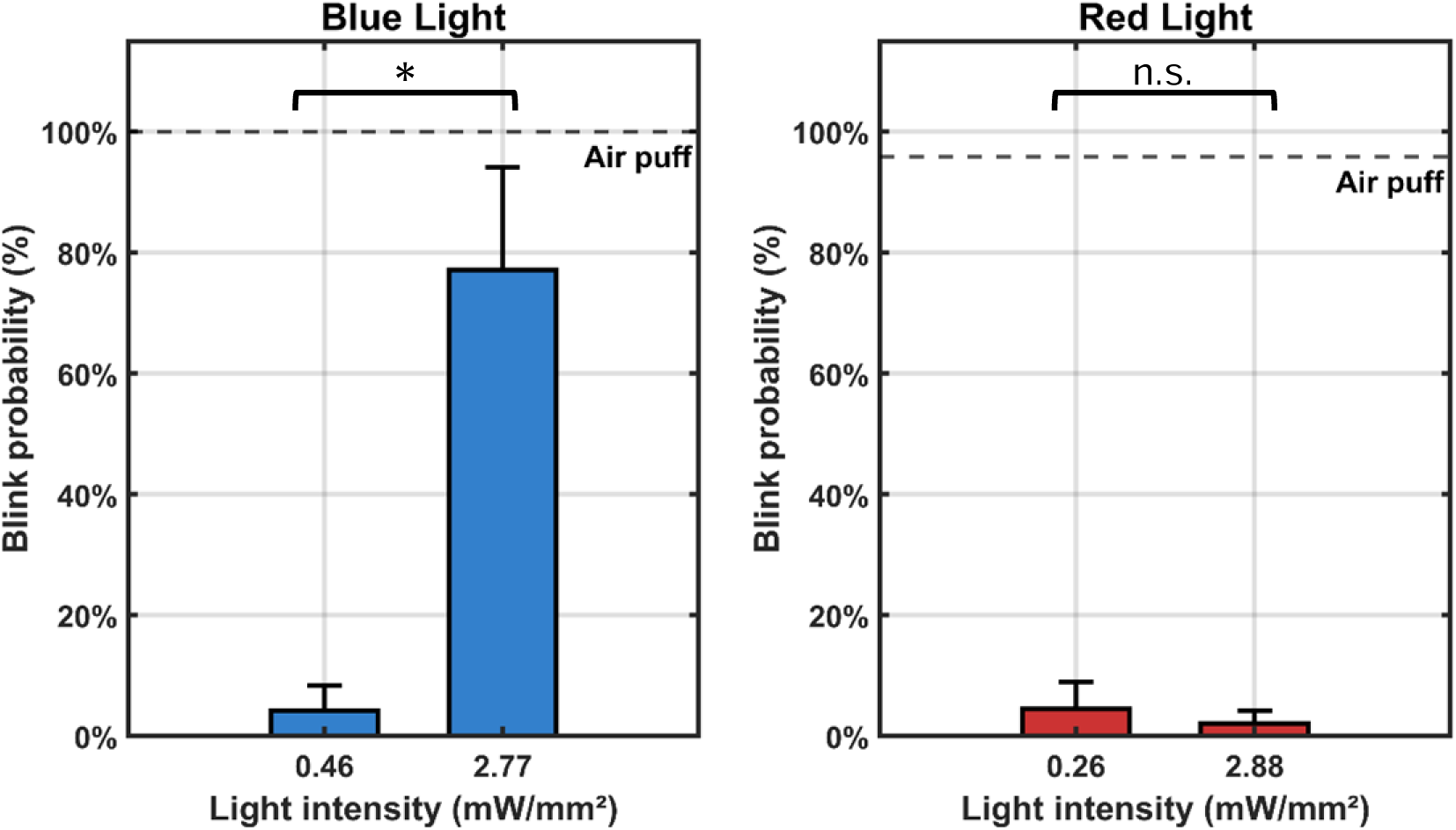
Blink probability depends on wavelength and intensity of light stimulation in TRPV1-ChR2-EYFP mice (N=3). Blink probability increased with light intensity under blue light stimulation, whereas no comparable intensity-dependent response was observed with red light stimulation (outside the excitation spectrum of channelrhodopsin). Data represent mean ± SEM across animals (p<0.05).

### 2.2. Machine learning-based pose estimation identifies multiple measurable features that characterize blink reflex

We employed machine learning-based toolkits to detect six previously-identified and validated features of reflexive blinking behavior, which were used to classify responses to corneal stimulation. Using the open-source pose estimation toolkit (DeepLabCut, or “DLC”, Genève, Switzerland), we trained custom transfer learning-based deep neural networks to detect eight anatomical landmarks along the mouse eye-eyelid boundary across video frames: [1] inner and [2] outer corners of the eye; centers of the [3] upper and [4] lower eyelids; and midpoints [5][6][7][8] of each quadrant (**Fig. 3A**). Using these eight planar coordinates, we subsequently calculated six metrics that characterized the blink reflex across each video frame: (1) Euclidean distance, or the distance between the centers of the upper and lower eyelids (**Fig. 3B**); (2) the eye aspect ratio, or the ratio between the averaged height and the length of the eye (**Fig. 3C**); (3) and (4) the eye angles of the inner and outer corners, respectively (**Fig. 3D**); (5) the total surface area of the eye (**Fig. 3E**); and (6) the velocity, or the change in Euclidean distance from the previous frame (**Fig. 3F**). We successfully calculated these across the frames for all 767 trials, including air puff and blue light stimuli. The best-performing custom DeepLabCut model, trained on 250 labeled frames, achieved an average root mean squared error (RMSE) of 1.00 pixels on the video dataset (***Supplementary Figure 1***).

**Figure 3.**
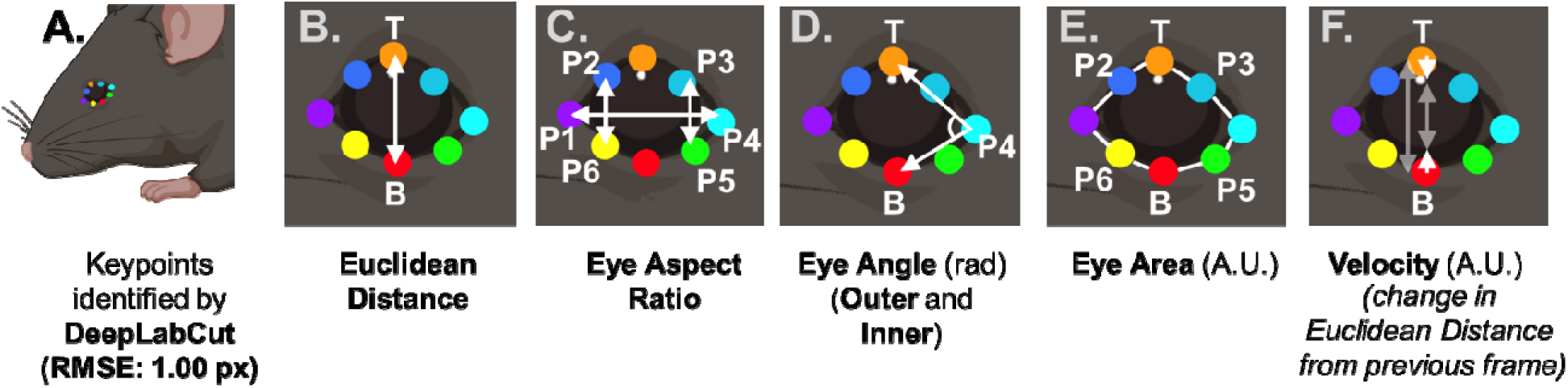
Calculation of six blink features from DeepLabCut-identified periocular key points. Features span from Euclidean distance, eye aspect ratio, outer and inner eye angles, eye area, and velocity (change in Euclidean distance from the previous frame).

To improve the robustness of our model, we trained the model not only on open-eye blinks but also on sensitized, partially closed eye-states to prevent boundary-tracking failure. Capsaicin or a vehicle control solution was applied prior to dedicated recording sessions to induce ocular sensitization associated with sustained squinting behavior. This manipulation altered the eye-eyelid boundary during recordings and was included to increase morphological variability during model development, thereby improving the robustness of the trained model to changes in eyelid position. Data acquired under these conditions were incorporated into model training and validation datasets and were not treated as a primary experimental condition in the main analyses.

### 2.3. Univariate binary classification via thresholding accurately detects evoked blink reflex following corneal stimulation

We tested the ability to automate the detection of blink reflex across hundreds of trials in an efficient manner using both simple binary thresholds and machine learning-based approaches. This enabled a comparison between univariate and multivariate classification of trials by the presence of blink behavior. We assessed each method for accuracy relative to a manually scored gold standard by two independent observers.

For univariate analysis, we used the optimal threshold value for each metric to perform binary classification of the recorded trials. For each feature characterizing the blink reflex, we plotted said metric across frames. If this value fell either above or below said threshold for a given metric (i.e., Euclidean distance, eye aspect ratio, eye angle), a given trial could be classified as either positive or not negative for blink. This threshold value was held constant across trials for each given metric and could be adjusted to assess its impact on the accuracy of binary classification and optimized for the highest accuracy (**Supplementary figure 1**). Once optimized, all six features demonstrated high accuracy (>92%) in classifying trials relative to a gold standard (manual scoring) (**Figure 4**). Five of six features exhibited higher accuracy (>98%) (**Figure 6**). Only two out of 768 trials were excluded due to ambiguity in blink response, even with manual observation. These accuracies indicate that most, if not all, metrics can accurately characterize the blink reflex following corneal stimulation.

**Figure 4.**
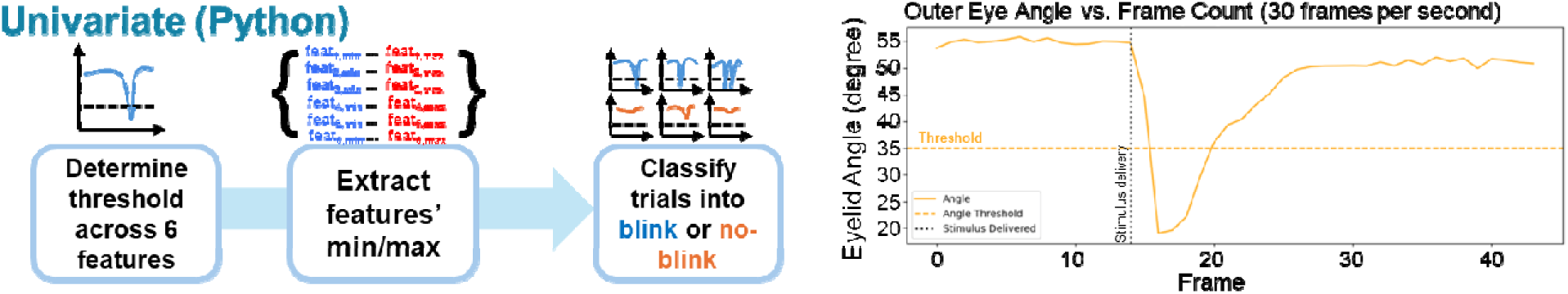
Threshold-based binary classification of evoked blink reflex from individual features. For each trial and blink reflex feature, the corresponding metric was plotted across video frames acquired at 30 frames per second. Trial-wise minimum and maximum values were extracted and compiled into a separate dataset. A global threshold, derived across all trials, was applied to classify trials for the presence of a blink reflex based on whether the extracted minima or maxima crossed the threshold, depending on the metric analyzed. This approach enabled rapid, high-throughput binary classification of blink responses. However, although global thresholding permits reliable detection, it does not precisely determine blink timing, as inferred onset varies with the selected threshold value.

### 2.4. Multivariate binary classification via machine learning algorithms accurately detects evoked blink reflex following corneal stimulation

We also utilized a multivariate approach comparing machine learning models in their ability to classify trials by blink reflex using the minima and maxima of the six metrics identified (Automated Machine Learning, or AutoML; Oracle AutoML).(Yakovlev et al., 2020) As these features were derived from the same eight key points, it was critical to reduce the number of collinear metrics, as multicollinearity undermines the predictive value of these models. Using standards for collinearity (<0.4), we pared down the total metric values from 12 to 4, based on the metric with the highest correlation (minimum outer eye angle), as well as three additional maxima (maximum eye aspect ratio; maximum velocity; and maximum eye area). Seven different models were tested, in order of complexity: Logistic regression classifier; Gaussian Naïve Bayes (GaussianNB); KNeighborsClassifier; Support Vector Classifier (SVC); Light Gradient Boosting Machine (LGBM) Classifier; XGBoost Classifier (XGBClassifier); and CatBoost Classifier. Based on balanced accuracy, we identified the tree-based Light Gradient-Boosting Machine (LGBM) classifier (98%): the near-perfect area under the Receiving Operating Characteristic (ROC) curve (or “AUC”) of 0.98 indicates high accuracy in classifying blink reflex, further verified by the LGBM classifier’s confusion matrix indicating low false positive (2 trials) and false negative counts (1 trial) within the testing set (N_total_=154 trials). However, multiple other algorithms exhibited similarly high accuracy, including the sigmoidal logistic regression classifier, the CatBoost classifier (decision-tree algorithm), and the random-forest XGBoost classifier. The complexity of the model did not predict the accuracy, as most models, independent of complexity, exhibited similar high accuracy above 90% (**Figure 5**). In aggregate, these accuracies suggest multiple features extracted from pose estimation can sufficiently characterize the blink reflex in a reproducible and reliable manner, whether singly or in aggregate.

**Figure 5.**
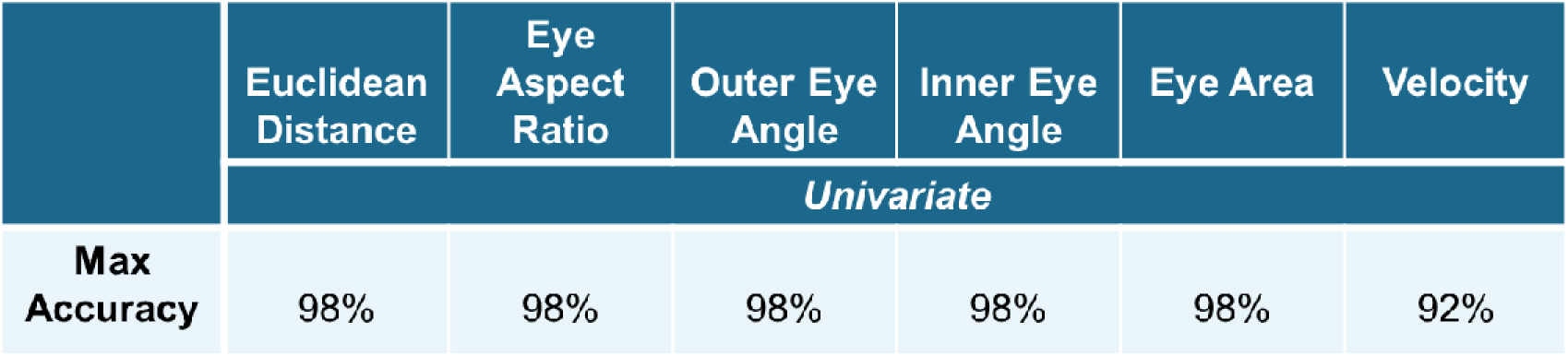
High classification accuracy via automated univariate thresholding of blink features. For all metrics, most trials could be classified for blink reflex with high accuracy (>92%) relative to manual assessment. Most metrics, except for velocity, exhibited greater accuracy than this minimum.

**Figure 6.**
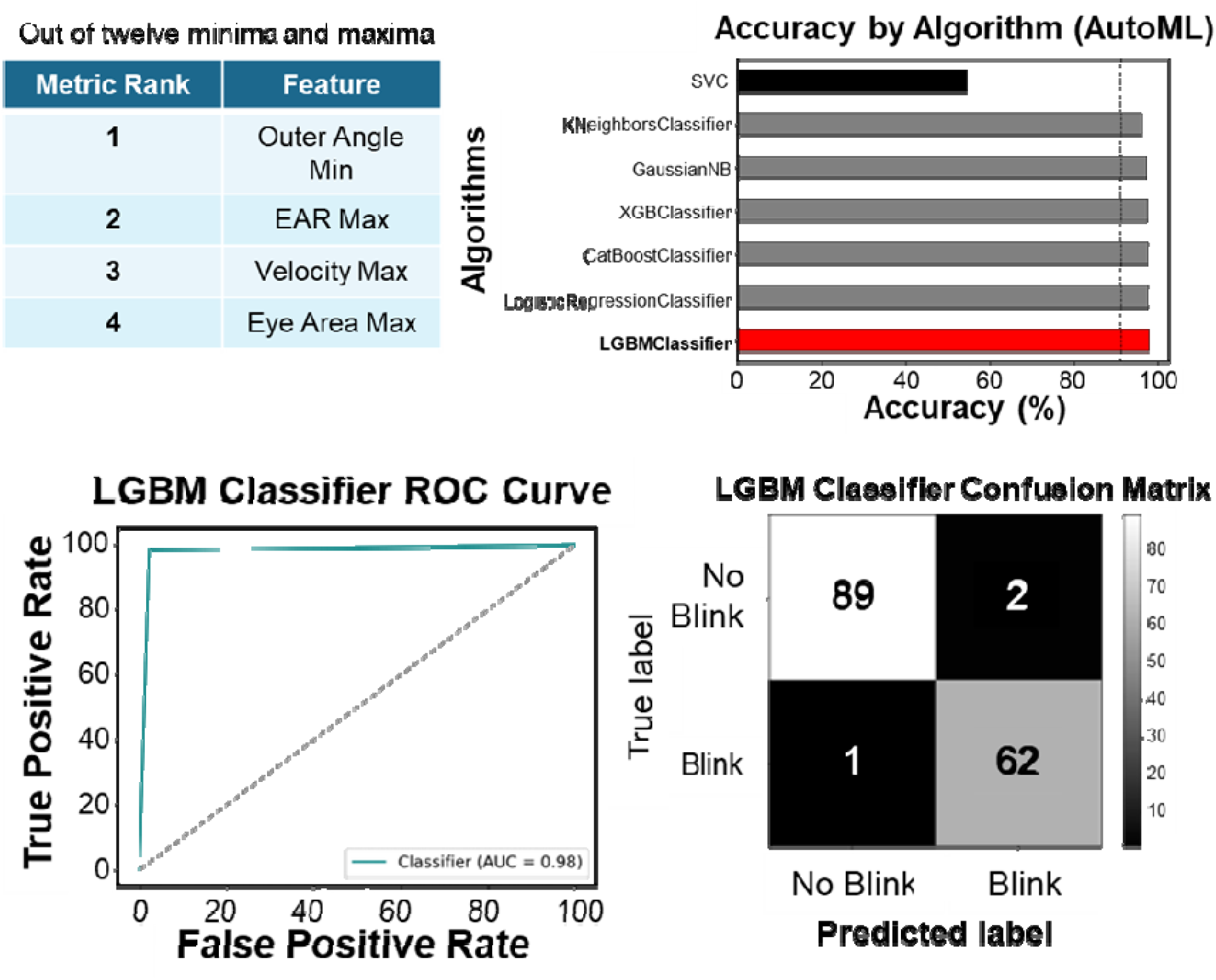
High-accuracy multivariate classification of blink response via AutoML. Based on a dataset of four values out of the twelve total minima and maxima for the six metrics, multiple models could perform binary classification of trials with high accuracy. Of these, the tree-based Light Gradient Boosting Machine classifier demonstrated the highest balanced accuracy.

## 3. Discussion

### 3.1. Optogenetic stimulation of polymodal corneal nociceptors reliably evokes blink reflexes in transgenic mice

In this report, we validate the use of optogenetic stimulation of corneal nociceptors and automate the detection of painful blink reflexes in mice. As demonstrated with both pressurized air and blue light, supra-threshold corneal stimulation reliably elicits reflexive behavioral responses, providing the consistency necessary for entraining operant behaviors (i.e., licking) that may serve as readouts of pain perception in future studies (Andermann, 2010; Black et al., 2023, 2020; Freund and Walker, 1972). We characterized the blink reflex across a range of intensities and with wavelengths of light inside and outside of the excitation range of channelrhodopsin. Mice exhibited robust blink responses to corneal stimulation with blue light, but not red light, suggesting optogenetic activation of these nociceptors above threshold intensity reproducibly induces blink behavior. This aligns with our prediction that optogenetic corneal stimulation in this transgenic model can be used to induce reflexive behavior. The consistency of the blink reflex to air puff (>95%) across sessions demonstrates that the transgenic mice studied here are consistent with the evoked behavior observed in prior literature with wildtype and genetically modified mice. TRPV1-ChR2-EYFP mice exhibit comparable rates of blink reflex across trials with optogenetic stimuli and air puff, suggesting that optogenetic stimulation engages ocular neural circuits that overlap with those recruited by noxious mechanical stimulation. Having characterized the blink reflex in a healthy state, we can now modulate this assay pharmacologically or chemically in future studies to examine the effects on blink reflex for well-established phenomena (i.e., sensitization via capsaicin). This also lays the foundation for assessing the blink reflex in models of disease state or genetic knockouts, assuming said mice can be crossbred with the TRPV1-ChR2-EYFP mouse line. Developing this assay represents a major milestone towards studying sub-second neural dynamics of blink reflex in healthy and disease states.

### 3.2. Automation of blink detection using machine learning

We have demonstrated that the blink reflex can be detected in a high throughput manner across hundreds of trials using either univariate or multivariate approaches based on six features extracted from recorded blink reflexes. This suggests that most features (with perhaps the exception of velocity) can reliably indicate blink reflex in each trial; for example, the change in eye angle indicates blink reflex with similar accuracy as the minimum Euclidean distance. Therefore, these features are redundant for detecting blink reflex (an advantage for experimental setups or models that may visually occlude portions of the eye or obscure certain features). Notably, multivariate classification did not improve blink detection accuracy compared to univariate classification. Detecting blink reflex validates the delivery of painful stimulation (both optogenetic and mechanical) activation of corneal nociceptors and the sufficiency of said stimulus intensity to evoke a behavioral response.

We observed that blink reflex responses were stable across minor and natural variations in the geometry of stimulation. Although certain applications are sensitive to the incident angle of illumination, several unique details regarding the anatomy of the ocular surface and the motion of the eye mitigate these effects in our model. The corneal epithelium is densely innervated via a network of superficial nerve terminals distributed across its curved surface, such that small changes in illumination angle primarily change the spatial footprint of stimulation without substantially altering the overall population of afferents recruited. The use of collimated light, which maintains the beam diameter independent of distance, further reduces the sensitivity to positioning. In addition, natural rotational movements of the eye introduce trial-by-trial variations in the illumination angle under awake conditions. Despite these sources of variability, blink responses remained consistent across trials as detected by our automated pipeline, supporting the robustness of this approach to natural variations in stimulus delivery. These findings indicate that targeted optogenetic innervation of polymodal corneal nociceptors is sufficient to evoke robust and measurable blink behavior.

### 3.3. Automated blink detection enables temporally precise future mapping of corneal nociceptive circuits

We characterized and classified the evoked blink reflex to validate corneal nociceptor activation with the precision required to infer underlying neural activity in future studies. Automation of blink detection is key for reducing observer bias in blink detection and improves consistency in detection across sessions. This consistency is essential for larger datasets, such as electrophysiological or calcium-imaging data, for which the signals are small in scale, noisy, and exhibit high variability; high throughput data is fundamental in pinpointing the conserved signals fundamental to encoding pain. Automated blink detection streamlines the process of validating that neural data recorded concurrent with the observation of blink reflex reflects evoked activity. These results show that our approach achieves superior genetic specificity in targeting corneal nociceptors, allowing us to map cell-specific neural circuits with high temporal precision relative to the originating stimuli.

### 3.4. Automated detection of evoked blink reflexes facilitates validation of corneal pain pathway innervation across a broad stimulus range, with advantages over other behavioral readouts such as eye wiping and grimace

Automated blink reflex offers more consistent validation of corneal nociception across a range of stimulus intensities and modalities than other innate behavioral responses, such as eye wiping, squinting, and grimace. While valuable under certain conditions, these operant behaviors do not require entrainment but do include a somatic or temporal component (i.e., paw movement or sustained eyelid position). Eye-wiping with a forepaw, which reflects attempted removal of irritants, has previously been characterized for corneal nociception, particularly as a platform for opioids and tricyclic antidepressants (TCA) (Aicher et al., 2015; Farazifard et al., 2005; Hegarty et al., 2017; Li et al., 2019). However, it is most commonly measured seconds after ocular stimulation as the number of wipes in a given duration, greatly reducing its ability to identify causal relationships to neural activity (Cho et al., 2019; Farazifard et al., 2005; Hegarty et al., 2017). Squinting, or sustained orbital tightening involving the facial muscles, can reflect the presence of an environmental or administered stressor and has also been automated to assess pain (McCutcheon et al., 2024; Rea et al., 2022, 2018; Sowers, 2022). Orbital tightening in particular constitutes a major component of the more comprehensive mouse grimace pain assessment, which also includes nose and cheek bulge, ear position, and whisker changes (Langford et al., 2010; Rea et al., 2022, 2018). Grimace has been used to assess spontaneous pain in acute states more so than chronic pain states (Langford et al., 2010; Whittaker et al., 2021). However, a significant increase in pain is required to elicit detectable changes in the mouse grimace scale (Deuis et al., 2017). Additionally, grimace scale exhibits considerable variability in assessment when manually scored, especially for different pain states (Hohlbaum et al., 2020). While automation can address issues of consistency across pain states and between scorers, the mismatch between the duration of a pain state and the expression of grimace in mice limits its utility for mapping evoked pain.

### 3.5. Optogenetic innervation provides cellular and temporal specificity for corneal stimulation, enabling more precise mapping of the corneal pain pathway than either conventional mechanical approaches or recently developed light-exposure models

Conventional animal models of corneal pain enable circuit mapping but lack the cellular specificity required to differentiate the effects of different stimulus modalities and their corresponding nociceptive fiber subtypes. These models most often rely on one of two mechanical methods of stimulation: the nylon fiber-based Cochet-Bonnet aesthesiometers that measure filament bending or air-based Belmonte’s gas esthesiometer that delivers an air puff, most commonly carbon dioxide (Belmonte et al., 1999; Chao et al., 2015; Crabtree et al., 2024; Eio et al., 2025; Nosch et al., 2024). These act in different ways: the fiber-based Cochet-Bonnet enables a semi-quantitative assessment of mechanical sensitivity, with limited impact of thermal or cold stimulation, whereas the gas aesthesiometer can also test chemical sensitivity. This has implications for the types of nociceptors recruited within each model: the Cochet-Bonnet monofilament activates polymodal and mechano-nociceptive fibers through mechanical contact, air-based aesthesiometers recruit these same populations along with cold-responsive nociceptors, provided the temperature of the gas is sufficiently low. Additionally, neither method exclusively activates either polymodal or mechano-nociceptive nociceptors, as neither of these conventional models can recruit one subgroup without concurrently activating the other. Only the optogenetic approach outlined here allows for the targeted excitation of specific nociceptor subtypes, providing a level of control unattainable with conventional sensory stimuli.

Unlike our approach, which uses light as a precise stimulus, existing blue light-exposure models focus on inducing hypersensitivity and chronic neural remodeling, similar to traditional surgical methods. These blue light-exposure models utilize long-term exposure across 12-hour blue light and dark cycles in wildtype C57BL/6 mice to induce hyperalgesia (Gao et al., 2023; Lee et al., 2016). This non-specific, low-grade exposure of blue light over multiple days and weeks in wildtype mice differs from the optogenetic approach laid out in this study, which delivers short (10 millisecond) high-intensity pulses of blue light to activate corneal nociceptors in transgenic mice to evoke acute pain. While both blue-light exposure models and our optogenetic approach utilize similar wavelengths, they differ immensely in the duration and the intensity of the light, contrasting broad environmental exposure with discrete, targeted application. Indeed, these light-exposure models operate closer to other environmental exposure models such as low-humidity models of dry eye disease (Barabino et al., 2005). Prolonged optogenetic activation of corneal nociceptors via blue light in transgenic TRPV1-ChR2-EYFP mice has also been used to study the role of corneal afferents in neuroinflammation (Yun et al., 2026). However, prior to this work, optogenetic activation of polymodal corneal nociceptors in this transgenic line had not been used to elicit nocifensive behavioral responses in awake mice.

### 3.6. Limitations and future directions

Our work focused on automated blink reflex detection associated with acute corneal nociception, which enables systematic mapping of corneal pain processing under a healthy state. We demonstrated the ability to automate the detection of blink reflex across hundreds of trials. Whereas these trials were not all fully independent, blink reflex was consistently evoked across animals, supporting the reliability of this behavior. Future studies with finer light intensity resolutions will be needed to determine whether nociceptor-specific optogenetic corneal stimulation produces a dose-dependent blink response (i.e. psychometric curve).

Pathological or chronic pain states may alter certain behavioral and morphological features measured here (i.e., squinting), which would require minimal calibration of our proposed framework to accommodate these changes. Here we began to characterize the blink reflex in the healthy state following the activation of TRPV1-expressing nociceptors. Future studies will be important to assess these behavioral responses following sensitization via chemical application and/or injury, as well as to establish the stability of baseline blink responses across repeated sessions within the same animals. We also emphasize that the automation of blink reflex alone is insufficient to validate corneal pain in mice. Rather, it provides a scalable behavioral indicator when used alongside complementary measures such as licking, for a comprehensive validation of operant and reflexive components of corneal pain in rodents. Follow-up studies can incorporate entrained responses, in addition to operant behaviors, to strengthen the behavioral evidence for corneal pain.

In this study, we used optogenetic activation of TRPV1-expressing polymodal corneal nociceptors, which respond to thermal and mechanical stimuli. While these represent a major functional class of nociceptors in the cornea, other populations – including mechano-nociceptors and cold-sensitive nociceptors – were not directly and specifically targeted or characterized (beyond mechanical stimulation provided via air puff). This paradigm and automated pipeline may be adaptable to these modalities using alternate promoters in transgenic mouse lines (e.g., TRPM8 or TRPM3).

Lastly, our work exclusively utilized male mice. Future studies will incorporate both male and female mice to improve the robustness of these findings across sexes and to assess potential sex-specific differences in nociceptive blink reflexes.

## 4. Conclusions

We have characterized blink reflex behaviors in response to corneal optogenetic stimulation in transgenic mice with improved nociceptive specificity and established an automated framework for high-throughput and observer-independent empirical assessment of blink response across large-scale datasets. Compared to conventional air puff paradigms, our optogenetic approach enables temporally-precise and cell-type-specific stimulation of corneal nociceptive afferents while supporting repeat acquisition of blink behavior across hundreds of trials. Our automated pipeline enabled consistent and objective assessment of large-scale video-based blink behavior and successful classification into blink or no-blink responses via both univariate and multivariate approaches. Together, these findings establish a framework for automated and TRPV1-specific behavioral assessment of corneal nociception that extends prior optogenetic paw-withdrawal paradigms for somatic pain in the same transgenic model and may support future studies of other corneal nociceptor populations. This framework can be extended to evaluate corneal pain using automated readouts, including integration with operant self-report paradigms such as licking, as demonstrated in somatic hindpaw pain in head-restrained mice (Black et al., 2020; Guo et al., 2014). This paradigm also provides a basis for future studies examining sub-second neural dynamics underlying blink behavior across the somatosensory cortex and related brain regions.

## Declaration of Generative AI and AI-assisted technologies in the writing process

During the preparation of this work, the authors used ChatGPT to aid in editing for style and rephrasing for clarity. After using these tools, the authors reviewed and edited the content as needed and take full responsibility for the content of the published article.

## References

Acosta, M.C., Tan, M.E., Belmonte, C., Gallar, J., 2001. Sensations evoked by selective mechanical, chemical, and thermal stimulation of the conjunctiva and cornea. Invest Ophthalmol Vis Sci 42, 2063–2067.

Adamantidis, A.R., Zhang, F., de Lecea, L., Deisseroth, K., 2014. Optogenetics: Opsins and Optical Interfaces in Neuroscience. Cold Spring Harbor Protocols 2014, pdb.top083329-pdb.top083329. 10.1101/pdb.top083329

Aicher, S.A., Hermes, S.M., Hegarty, D.M., 2015. Denervation of the Lacrimal Gland Leads to Corneal Hypoalgesia in a Novel Rat Model of Aqueous Dry Eye Disease. Invest. Ophthalmol. Vis. Sci. 56, 6981. 10.1167/iovs.15-17497

Andermann, M.L., 2010. Chronic cellular imaging of mouse visual cortex during operant behavior and passive viewing. Front. Cell. Neurosci. 10.3389/fncel.2010.00003

Barabino, S., Shen, L., Chen, L., Rashid, S., Rolando, M., Dana, M.R., 2005. The Controlled-Environment Chamber: A New Mouse Model of Dry Eye. Invest. Ophthalmol. Vis. Sci. 46, 2766. 10.1167/iovs.04-1326

Belmonte, C., Acosta, M.C., Schmelz, M., Gallar, J., 1999. Measurement of corneal sensitivity to mechanical and chemical stimulation with a CO2 esthesiometer. Invest Ophthalmol Vis Sci 40, 513–519.

Black, C.J., Allawala, A.B., Bloye, K., Vanent, K.N., Edhi, M.M., Saab, C.Y., Borton, D.A., 2020. Automated and rapid self-report of nociception in transgenic mice. Scientific Reports 10. 10.1038/s41598-020-70028-8

Black, C.J., Saab, C.Y., Borton, D.A., 2023. Transient gamma events delineate somatosensory modality in S1. 10.1101/2023.03.30.534945

Boyden, E., 2011. A history of optogenetics: the development of tools for controlling brain circuits with light. F1000 Biology Reports 3. 10.3410/B3-11

Chao, C., Stapleton, F., Badarudin, E., Golebiowski, B., 2015. Ocular Surface Sensitivity Repeatability with Cochet-Bonnet Esthesiometer. Optometry and Vision Science 92, 183–189. 10.1097/OPX.0000000000000472

Cho, J., Bell, N., Botzet, G., Vora, P., Fowler, B.J., Donahue, R., Bush, H., Taylor, B.K., Albuquerque, R.J.C., 2019. Latent Sensitization in a Mouse Model of Ocular Neuropathic Pain. Trans. Vis. Sci. Tech. 8, 6. 10.1167/tvst.8.2.6

Crabtree, J.R., Tannir, S., Tran, K., Boente, C.S., Ali, A., Borschel, G.H., 2024. Corneal Nerve Assessment by Aesthesiometry: History, Advancements, and Future Directions. Vision 8, 34. 10.3390/vision8020034

Deisseroth, K., 2015. Optogenetics: 10 years of microbial opsins in neuroscience. Nature Neuroscience 18, 1213–1225. 10.1038/nn.4091

Deuis, J.R., Dvorakova, L.S., Vetter, I., 2017. Methods Used to Evaluate Pain Behaviors in Rodents. Frontiers in Molecular Neuroscience 10. 10.3389/fnmol.2017.00284

Eio, E., Yu, M., Liu, C., Lee, I.X.Y., Wong, R.K.T., Wong, J.H.F., Liu, Y.-C., 2025. Evaluation of Corneal Sensitivity: Tools We Have. Diagnostics 15, 1785. 10.3390/diagnostics15141785

Farazifard, R., Safarpour, F., Sheibani, V., Javan, M., 2005. Eye-wiping test: A sensitive animal model for acute trigeminal pain studies. Brain Research Protocols 16, 44–49. 10.1016/j.brainresprot.2005.10.003

Fernández-Trillo, J., Florez-Paz, D., Íñigo-Portugués, A., González-González, O., Del Campo, A.G., González, A., Viana, F., Belmonte, C., Gomis, A., 2020. Piezo2 Mediates Low-Threshold Mechanically Evoked Pain in the Cornea. J. Neurosci. 40, 8976–8993. 10.1523/JNEUROSCI.0247-20.2020

Freund, G., Walker, D.W., 1972. Operant conditioning in mice. Life Sciences 11, 905–914. 10.1016/0024-3205(72)90042-2

Gao, N., Lee, P.S.Y., Zhang, J., Yu, F.X., 2023. Ocular nociception and neuropathic pain initiated by blue light stress in C57BL/6J mice. Pain 164, 1616–1626. 10.1097/j.pain.0000000000002896

Gharagozloo, M., Amrani, A., Wittingstall, K., Hamilton-Wright, A., Gris, D., 2021. Machine Learning in Modeling of Mouse Behavior. Front. Neurosci. 15, 700253. 10.3389/fnins.2021.700253

Gong, X., Mendoza-Halliday, D., Ting, J.T., Kaiser, T., Sun, X., Bastos, A.M., Wimmer, R.D., Guo, B., Chen, Q., Zhou, Y., Pruner, M., Wu, C.W.-H., Park, D., Deisseroth, K., Barak, B., Boyden, E.S., Miller, E.K., Halassa, M.M., Fu, Z., Bi, G., Desimone, R., Feng, G., 2020. An Ultra-Sensitive Step-Function Opsin for Minimally Invasive Optogenetic Stimulation in Mice and Macaques. Neuron 107, 38–51.e8. 10.1016/j.neuron.2020.03.032

Guo, Z.V., Hires, S.A., Li, N., O’Connor, D.H., Komiyama, T., Ophir, E., Huber, D., Bonardi, C., Morandell, K., Gutnisky, D., Peron, S., Xu, N., Cox, J., Svoboda, K., 2014. Procedures for Behavioral Experiments in Head-Fixed Mice. PLoS ONE 9, e88678. 10.1371/journal.pone.0088678

Hegarty, D.M., Hermes, S.M., Yang, K., Aicher, S.A., 2017. Select noxious stimuli induce changes on corneal nerve morphology. J of Comparative Neurology 525, 2019–2031. 10.1002/cne.24191

Hohlbaum, K., Corte, G.M., Humpenöder, M., Merle, R., Thöne-Reineke, C., 2020. Reliability of the Mouse Grimace Scale in C57BL/6JRj Mice. Animals 10, 1648. 10.3390/ani10091648

Jeong, K. Figures created in BioRender.

Krzyzanowska, A., Pittolo, S., Cabrerizo, M., Sánchez-López, J., Krishnasamy, S., Venero, C., Avendaño, C., 2011. Assessing nociceptive sensitivity in mouse models of inflammatory and neuropathic trigeminal pain. Journal of Neuroscience Methods 201, 46–54. 10.1016/j.jneumeth.2011.07.006

Langford, D.J., Bailey, A.L., Chanda, M.L., Clarke, S.E., Drummond, T.E., Echols, S., Glick, S., Ingrao, J., Klassen-Ross, T., LaCroix-Fralish, M.L., Matsumiya, L., Sorge, R.E., Sotocinal, S.G., Tabaka, J.M., Wong, D., Van Den Maagdenberg, A.M.J.M., Ferrari, M.D., Craig, K.D., Mogil, J.S., 2010. Coding of facial expressions of pain in the laboratory mouse. Nat Methods 7, 447–449. 10.1038/nmeth.1455

Lee, H.S., Cui, L., Li, Y., Choi, J.S., Choi, J.-H., Li, Z., Kim, G.E., Choi, W., Yoon, K.C., 2016. Influence of Light Emitting Diode-Derived Blue Light Overexposure on Mouse Ocular Surface. PLoS ONE 11, e0161041. 10.1371/journal.pone.0161041

Li, F., Yang, W., Jiang, H., Guo, C., Huang, A.J.W., Hu, H., Liu, Q., 2019. TRPV1 activity and substance P release are required for corneal cold nociception. Nat Commun 10, 5678. 10.1038/s41467-019-13536-0

Li, T., Severson, K.S., Wang, F., Dunn, T.W., 2023. Improved 3D Markerless Mouse Pose Estimation Using Temporal Semi-supervision. Int J Comput Vis 131, 1389–1405. 10.1007/s11263-023-01756-3

Lin, J.Y., 2011. A user’s guide to channelrhodopsin variants: features, limitations and future developments: A user’s guide to channelrhodopsin variants. Experimental Physiology 96, 19–25. 10.1113/expphysiol.2009.051961

Marks, M., Qiuhan, J., Sturman, O., Von Ziegler, L., Kollmorgen, S., Von Der Behrens, W., Mante, V., Bohacek, J., Yanik, M.F., 2020. Deep-learning based identification, tracking, pose estimation, and behavior classification of interacting primates and mice in complex environments. 10.1101/2020.10.26.355115

Mattis, J., Tye, K.M., Ferenczi, E.A., Ramakrishnan, C., O’Shea, D.J., Prakash, R., Gunaydin, L.A., Hyun, M., Fenno, L.E., Gradinaru, V., Yizhar, O., Deisseroth, K., 2012. Principles for applying optogenetic tools derived from direct comparative analysis of microbial opsins. Nat Methods 9, 159–172. 10.1038/nmeth.1808

McCutcheon, N., Johnson, M.S., Rea, B., Ghumman, M., Sowers, L., Hultman, R., 2024. An Automated Squint Method for Time-syncing Behavior and Brain Dynamics in Mouse Pain Studies. JoVE 67136. 10.3791/67136

Mungalsingh, M.A., Thompson, B., Peterson, S.D., Murphy, P.J., 2025. Modelling the thermal effects of stimulus airflow from the Dolphin aesthesiometer on a model eye surface. Ophthalmic Physiologic Optic 45, 361–371. 10.1111/opo.13436

Nosch, D.S., Käser, E., Bracher, T., Joos, R.E., 2024. Clinical application of the Swiss Liquid Jet Aesthesiometer for corneal sensitivity measurement. Clinical and Experimental Optometry 107, 14–22. 10.1080/08164622.2023.2191782

Rea, B.J., Davison, A., Ketcha, M.-J., Smith, K.J., Fairbanks, A.M., Wattiez, A.-S., Poolman, P., Kardon, R.H., Russo, A.F., Sowers, L.P., 2022. Automated detection of squint as a sensitive assay of sex-dependent calcitonin gene–related peptide and amylin-induced pain in mice. Pain 163, 1511–1519. 10.1097/j.pain.0000000000002537

Rea, B.J., Wattiez, A.-S., Waite, J.S., Castonguay, W.C., Schmidt, C.M., Fairbanks, A.M., Robertson, B.R., Brown, C.J., Mason, B.N., Moldovan-Loomis, M.-C., Garcia-Martinez, L.F., Poolman, P., Ledolter, J., Kardon, R.H., Sowers, L.P., Russo, A.F., 2018. Peripherally administered calcitonin gene–related peptide induces spontaneous pain in mice: implications for migraine. Pain 159, 2306–2317. 10.1097/j.pain.0000000000001337

Sheppard, K., Gardin, J., Sabnis, G.S., Peer, A., Darrell, M., Deats, S., Geuther, B., Lutz, C.M., Kumar, V., 2022. Stride-level analysis of mouse open field behavior using deep-learning-based pose estimation. Cell Reports 38, 110231. 10.1016/j.celrep.2021.110231

Sowers, L.P., 2022. CGRP Administration Into the Cerebellum Evokes Light Aversion, Tactile Hypersensitivity, and Nociceptive Squint in Mice. Frontiers in Pain Research 3.

Spierer, O., Felix, E.R., McClellan, A.L., Parel, J.M., Gonzalez, A., Feuer, W.J., Sarantopoulos, C.D., Levitt, R.C., Ehrmann, K., Galor, A., 2016. Corneal Mechanical Thresholds Negatively Associate With Dry Eye and Ocular Pain Symptoms. Invest. Ophthalmol. Vis. Sci. 57, 617. 10.1167/iovs.15-18133

Varolgünes, N., Çelebisoy, N., Akyürekli, Ö., Pehlivan, M., Akyürekli, O., 1999. Analysis of the Corneal Reflex With Air Puff: Normal Controls and Patient Groups. Journal of Clinical Neurophysiology 16.

Whittaker, A.L., Liu, Y., Barker, T.H., 2021. Methods Used and Application of the Mouse Grimace Scale in Biomedical Research 10 Years on: A Scoping Review. Animals 11, 673. 10.3390/ani11030673

Yakovlev, A., Moghadam, H.F., Moharrer, A., Cai, J., Chavoshi, N., Varadarajan, V., Agrawal, S.R., Idicula, S., Karnagel, T., Jinturkar, S., Agarwal, N., 2020. Oracle AutoML: a fast and predictive AutoML pipeline. Proc. VLDB Endow. 13, 3166–3180. 10.14778/3415478.3415542

Yun, H., Moghbeli, K., Gerges, P.H., Cisney, R.N., Yoshida, M., Hawse, W.F., MacDonald, W.A., Bhuiyan, S.A., Renthal, W., Sullivan, C.J., Das, J., Kaplan, D.H., Singh, H., Davis, B.M., St. Leger, A.J., 2026. The production of the chemokine CCL2 by corneal sensory neurons initiates anti-viral immunity at the cornea and trigeminal ganglion. Cell Reports 45, 116693. 10.1016/j.celrep.2025.116693

