## Supplemental Figure 1 for "Automated detection of blink reflexes evoked by optogenetic stimulation of TRPV1-expressing corneal nociceptors in transgenic mice"

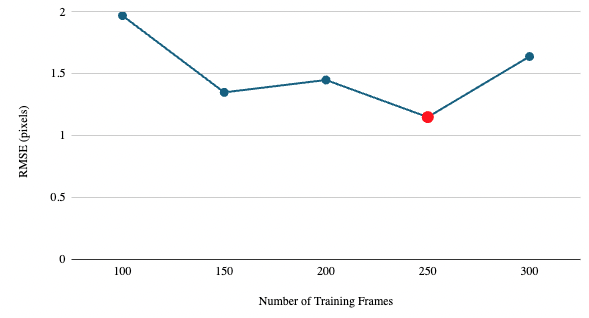

***Supplementary Figure 1. DeepLabCut model trained on 250 labeled frames produced the lowest root mean squared error (RMSE) between manual labels and predicted labels.*** *Models were trained using 100-300 labeled frames. Model performance was evaluated using root mean squared error (RMSE; in pixels), defined as the difference between manual annotated and model-predicted coordinates. The model trained on 250 labeled frames achieved the lowest RMSE and was selected for subsequent analysis.*
