## Supplemental Figure 2 for "Automated detection of blink reflexes evoked by optogenetic stimulation of TRPV1-expressing corneal nociceptors in transgenic mice"

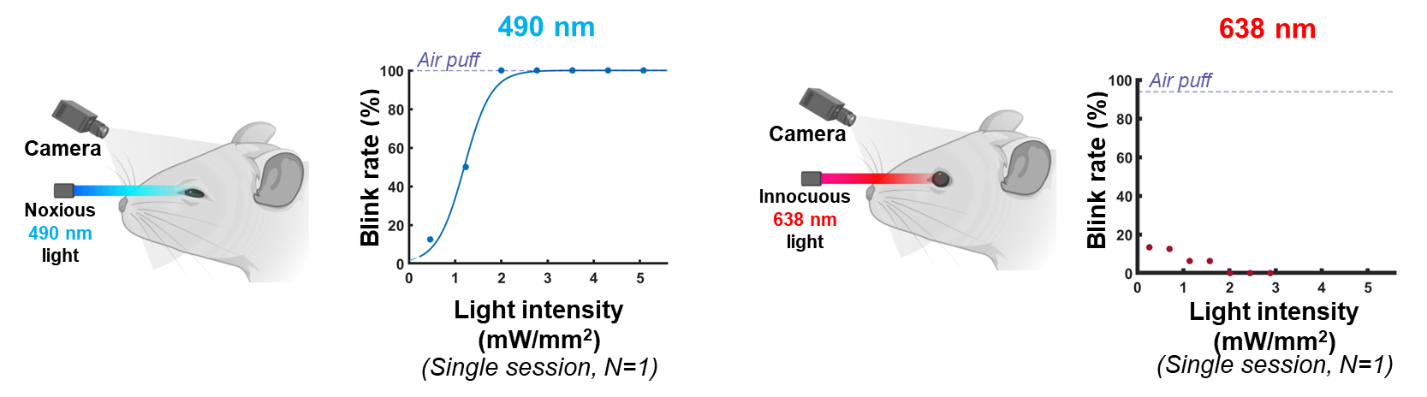

***Supplementary Figure 2. Blue light evokes graded blink responses in TRPV1-ChR2-EYFP mice.*** Data represents a single representative session from one animal in response to either optogenetic or control light, respectively. Light intensity (reported in mW/mm^2^) reflects the intensity of the stimulus delivered to the cornea. Blink rate (calculated across repeated trials at a given light intensity) rose from minimum response in a graded manner and plateaued at a maximum rate depending on the intensity of the blue light delivered**.** This graded response was not observed following the application of control red light (which falls outside the excitation range of channelrhodopsin). Sample single-session data demonstrate potential “dose-response” relationship between optogenetic corneal stimulation and blink reflex.
