## Supplemental Figure 3 for "Automated detection of blink reflexes evoked by optogenetic stimulation of TRPV1-expressing corneal nociceptors in transgenic mice"

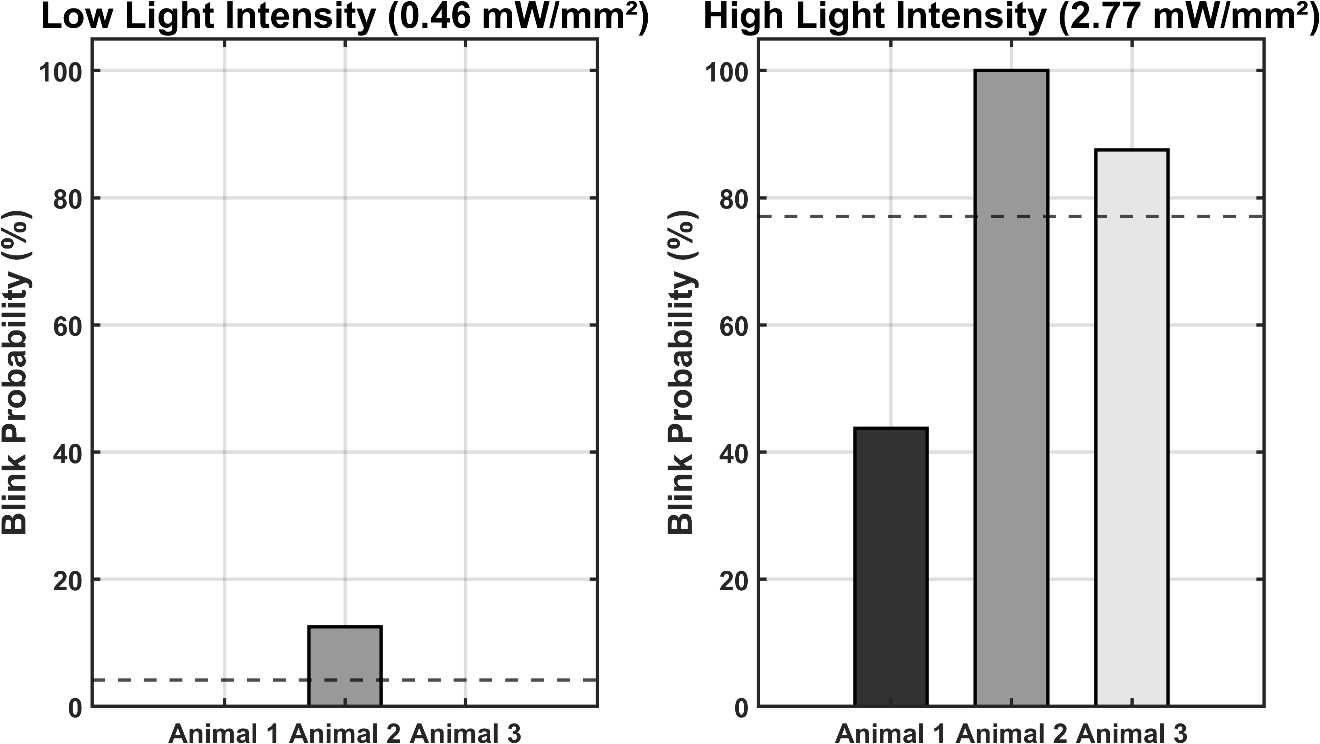

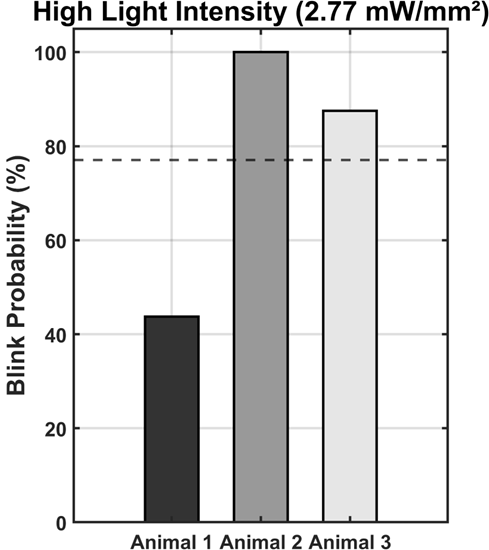

**Supplemental Figure 3. Blink responses by TRPV1-ChR2-EYFP individual mice.** Mice demonstrated near-zero blink response probability at low light intensity and increased blink probability at high light intensity, consistent with the averaged population response (dotted line, see Figure 2).
