## Supplemental Figure 4 for "Automated detection of blink reflexes evoked by optogenetic stimulation of TRPV1-expressing corneal nociceptors in transgenic mice"

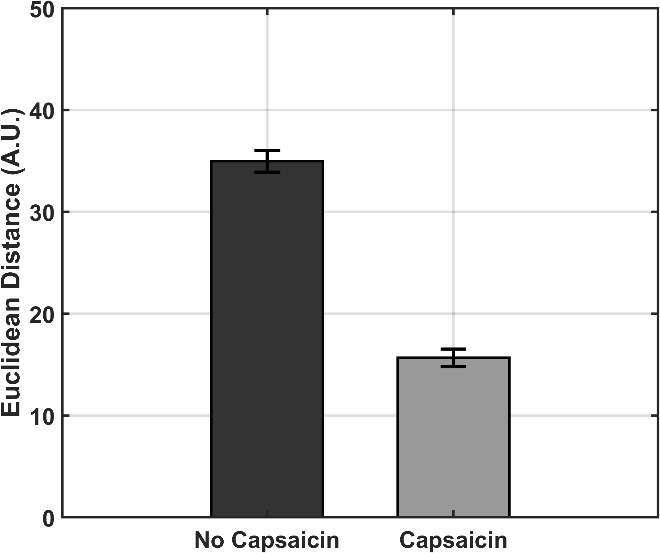

*p<0.01*

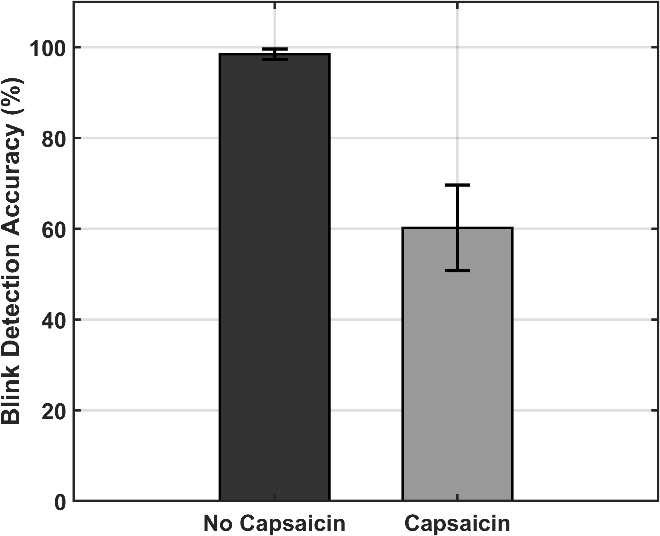

*p=0.06*

**Supplemental Figure 4. Capsaicin-induced morphological changes (i.e., squinting) alter blink response feature and blink detection accuracy.** Sessions involving capsaicin-induced squinting exhibited significantly reduced Euclidean distance between the upper to lower eyelid, consistent with decreased eye opening during squinting. Automated blink detection accuracy was also lower during capsaicin-associated squinting sessions (98% without capsaicin versus 60% with capsaicin), suggesting the need for classifier re-training under altered eye-geometry conditions in future studies involving sensitization or other forms of eye disease. Note that data from capsaicin-induced squinting were not included in the training of our machine learning models presented in Figures 5 and 6.
